# Efficient algorithms for designing maximally sized orthogonal DNA sequence libraries

**DOI:** 10.1101/2022.07.11.499592

**Authors:** Gokul Gowri, Kuanwei Sheng, Peng Yin

## Abstract

Orthogonal sequence library design is an essential task in bioengineering. Typical design approaches scale quadratically in the size of the candidate sequence space. As such, exhaustive searches of sequence space to maximize library size are computationally intractable with existing methods. Here, we present SeqWalk, a time and memory efficient method for designing maximally-sized orthogonal sequence libraries using the sequence symmetry minimization heuristic. SeqWalk encodes sequence design constraints in a de Bruijn graph representation of sequence space, enabling the application of efficient graph traversal techniques to the problem of orthogonal DNA sequence design. We demonstrate the scalability of SeqWalk by designing a provably maximal set of > 10^6^ orthogonal 25nt sequences in less than 20 seconds on a single standard CPU core. We additionally derive fundamental bounds on orthogonal sequence library size under a variety of design constraints.

## Introduction

Orthogonal DNA sequence libraries are sets of DNA sequences designed to have minimal crosstalk, which we broadly define as interaction with “off-target” sequences (Supplementary Note 1). In DNA-based biotechnologies, orthogonal sequences are widely used as cellular or molecular identifiers. For example, orthogonal sequences are used to barcode protein targets in DNA-based bioimaging (1), to label RNA molecules in individual cells for single-cell studies (2), and to program the assembly of components in a synthesis process (3), among many other applications (4–7).

The number of addressable features in these methods is dependent on the size of the orthogonal DNA sequence library that is used. For example, the multiplexity of DNA-based multiplexed epitope imaging is constrained by the number of orthogonal barcode sequences (1). Despite this, orthogonal DNA sequence library design methods are typically ad hoc, and do not yield optimal or maximally-sized sets of sequences. This can ultimately limit the scalability of the intended experimental application.

Existing methods for orthogonal DNA sequence library design typically use the following approach: First, a set of *S* candidate sequences is selected. Then, each pair of sequences in the candidate pool is compared to estimate crosstalk, resulting in an *S* × *S* crosstalk matrix which is used to select a subset of sequences with the desired degree of orthogonality (4, 6, 8). This approach requires a number of pairwise comparisons that scales quadratically in the number of candidate sequences, and becomes practically infeasible when the candidate sequence space is large. For example, screening the entire space of possible 25mers would require ~ 10^30^ crosstalk comparisons, which even at 1 nanosecond per comparison would take over 10^5^ times the age of the known universe. As a result, existing methods for large orthogonal sequence library design search only a small portion of sequence space (4, 6), precluding the design of maximally-sized libraries. Furthermore, large library design tasks have required high performance computing resources (6).

Sequence-based heuristics, such as sequence symmetry minimization (SSM), can be used to predict the orthogonality of a set of sequences. A set of sequences is considered to satisfy SSM for length *k* if no subsequence of length *k* appears more than one time in the set. SSM and closely related heuristics have been widely used for the design of orthogonal sequence libraries, with extensive experimental validation in several application contexts (3, 6, 9, 10).

In this work we develop a scalable computational tool which enables the design of maximal orthogonal sequence libraries that minimize fundamental limits on experimental methods. We use a de Bruijn graph representation of SSM constraints, which allows the application of efficient and theoretically tractable algorithms. Using our graph-theoretic approach, we design orthogonal 25nt sequence libraries of unprecedented size (> 10^6^ sequences) in less than one minute on a single standard CPU core, with provable guarantees of maximality and predicted orthogonality using SSM. The tools and theory presented in this work can be used to guide the design of DNA-barcoded molecular systems and to maximize the scalability of DNA engineering-based biotechnologies.

## Results

### k-mer graphs for orthogonal sequence design

We have developed SeqWalk, a computational tool for designing maximal orthogonal sequence libraries through the application of efficient graph-based algorithms.

The key observation underlying SeqWalk is that orthogonality constraints in sequence design problems can be naturally encoded in de Bruijn graph representations of sequence space. De Bruijn graphs, also known as k-mer graphs, are sequence representations that have been well studied in discrete mathematics (11–13). A k-mer is a length *k* sequence. A k-mer graph has all possible k-mers as nodes, and edges between k-mers that overlap by *k* – 1 symbols. In particular, if a k-mer *k*_1_ can be transformed into a k-mer *k*_2_ by removing its first symbol and appending a symbol, then there is a directed edge from *k*_1_ to *k*_2_.

On a k-mer graph, a length *L* sequence can be represented as a path over *L* – *k* + 1 nodes. The traversed nodes will correspond to each k-mer that appears in the sequence. A set of sequences that can be represented as non-intersecting paths on a k-mer graph share no common k-mers, and thus satisfy SSM for the corresponding *k*. This points toward a method for generating sequences that implicitly satisfy SSM for length *k*: one can simply select several non-intersecting paths on a k-mer graph. One way to produce non-intersecting paths on a graph is to take a single self-avoiding walk, and then partition this walk into multiple non-intersecting paths. The longest possible self-avoiding walks on a graph are Hamiltonian paths, which visit every node of the graph exactly one time. A partitioned Hamiltonian path will result in sequences that fully occupy k-mer space, and thus yield maximally sized orthogonal sequence libraries.

In SeqWalk, we apply a recently discovered mathematical technique for traversing de Bruijn graphs, which yields Hamiltonian paths in *O*(1) time and memory per node (13), to efficiently and scalably design orthogonal sequence libraries. The main SeqWalk algorithm is remarkably simple: our implementation requires less than 100 lines of code, including output formatting (Supplemental File 1).

### Performance benchmarks

To understand the practical relevance of the high efficiency of SeqWalk, we perform benchmark analysis against a traditional pairwise comparison approach for designing SSM satisfying sequence libraries, which is theoretically equivalent to the network elimination algorithms used in existing orthogonal sequence design methods (Supplemental File 2) (4, 6). We find that SeqWalk produces a larger number of sequences in less time than convergence of the pairwise comparison method for every tested design problem, with performance gains empirically increasing for design tasks of increasing complexity (Fig. 2). In the case of SSM *k* = 8 for 20nt sequences, SeqWalk produces more than 10 times as many orthogonal sequences as the pairwise algorithm, in approximately 1% of the computation time (Supplemental File 2).

**Fig. 1.**
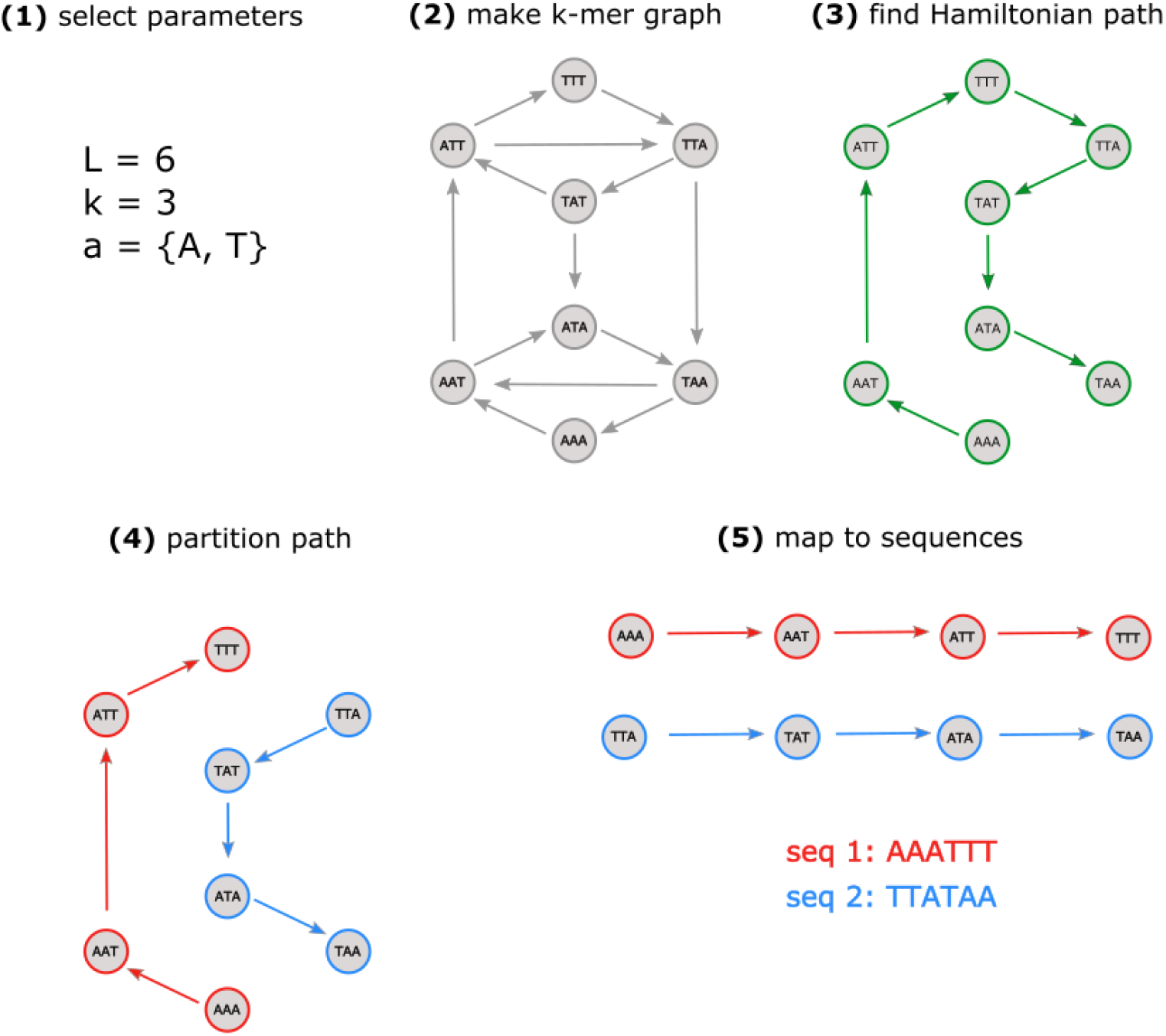
Workflow of graph-based sequence design algorithm. **(1)** Select length of desired sequences *L*, length of prevented substrings (SSM constraint) *k*, and allowed alphabet *a.* **(2)** Represent sequence space using a k-mer graph. **(3)** Select a random Hamiltonian path in k-mer graph. **(4)** Partition the selected Hamiltonian path into fragments with *L* – *k* +1 nodes. **(5)** Map fragments to corresponding sequences.

**Fig. 2.**
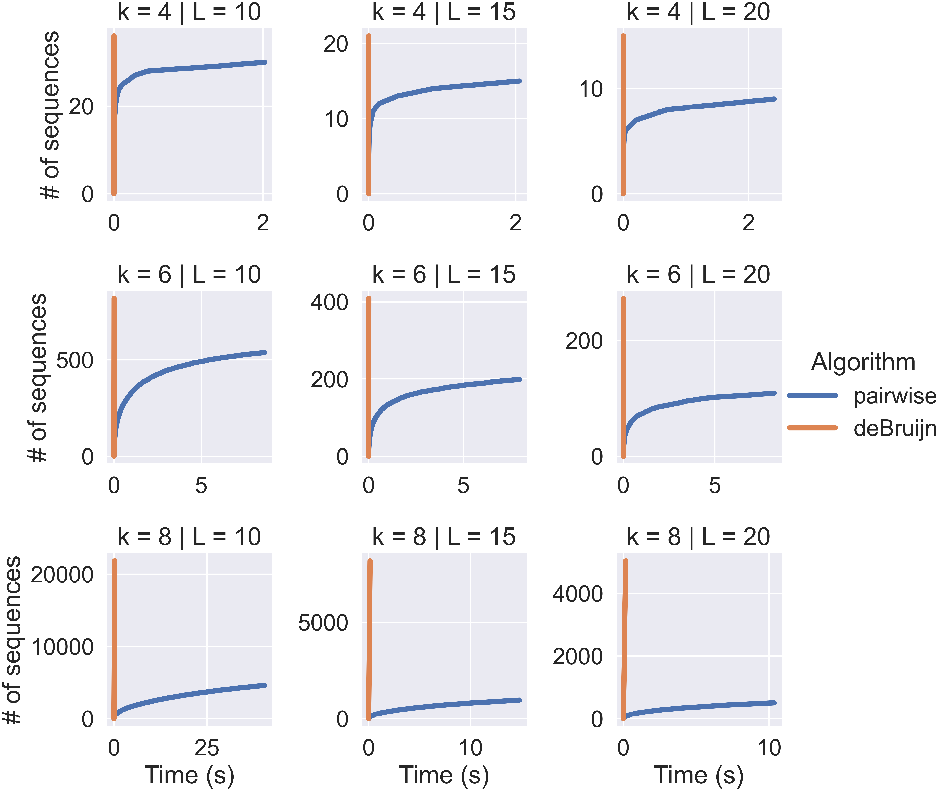
Performance of graph-based sequence design method (orange) in comparison to method involving pairwise crosstalk evaluations (blue), for several design problems. Each plot shows trajectories of the number of sequences designed in wall clock time for each method, averaged over 10 trials (see Supplement for details). Plots on the same row have the same *k* value (length of prevented substrings), and plots on the same column have the same *L* value (sequence length). All plots are for the case of a 4 letter alphabet.

**Fig. 3.**
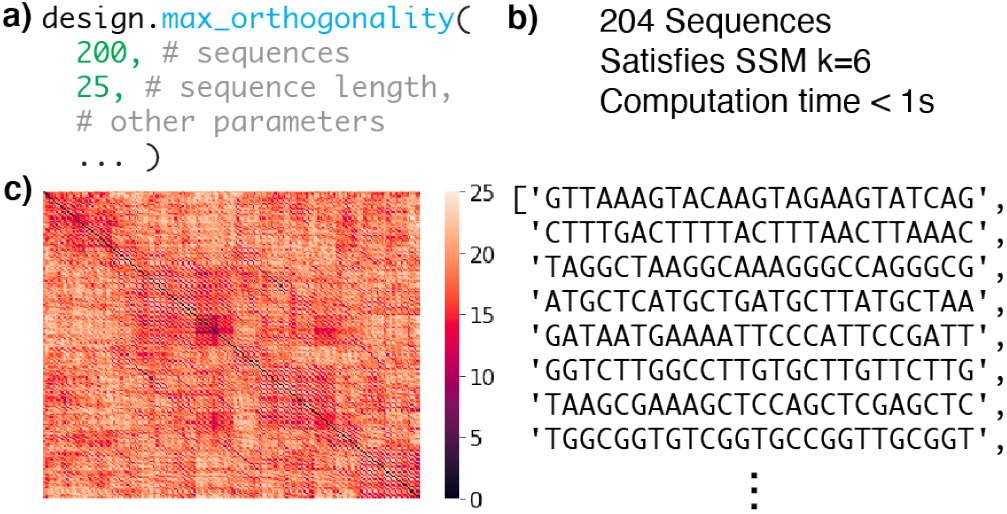
Depiction of seqwalk software package. **(a)** Example design code which produces a library of at least 200 25nt sequences with maximal orthogonality according to the SSM heuristic. **(b)** Output of example design code. **(c)** Crosstalk analysis of designed library, using Hamming distance. Each row/column represents a sequence, and each entry is colored by Hamming distance.

The largest published library of orthogonal sequences consists of approximately 2.4 * 10^5^ 25nt sequences satisfying SSM for *k* = 12. The library was designed with a pairwise sequence comparison algorithm, and used high performance computing tools (6). Using SeqWalk, we are able to generate over 1.2 * 10^6^ orthogonal 25nt DNA sequences satisfying SSM for *k =* 12, in less than 17 seconds on a single CPU core (Supplemental File 1). While previous methods subsample sequence space for candidate sequences (4, 6), SeqWalk exhaustively traverses sequence space, with a candidate sequence pool 9 orders of magnitude larger than previous work.

### Sequence design under additional constraints

In many applications, there are additional constraints to orthogonal sequence libraries beyond limitations on off-target binding. One common constraint is the prevention of crosstalk with reverse complements of sequences in the library. For sequence design under this constraint, SeqWalk integrates two efficient algorithms: a filtering-based method for 3-letter libraries, and an adaptation of the Hierholzer algorithm (14) for 4-letter libraries (Supplementary note 4).

SeqWalk design can also consider other common constraints such as requiring GC content within a window, absence of specific sequence patterns, and the absence of significant secondary structure. We provide efficient algorithms for filtering SeqWalk libraries for these characteristics (Supplementary notes 2, 3, 4, 5). We find that 3-letter SeqWalk libraries are particularly amenable to such filtering, as they have sequences with lower variance in GC content (Fig. S4), low prevalence of secondary structure (Fig. S5) and little crosstalk with reverse complements (Supplementary note 4).

### Theoretical results

Typical approaches to orthogonal sequence design provide no theoretical guarantees of optimality. The theoretical tractability of SeqWalk allows us to design sequence libraries that are provably maximally sized or provably maximally orthogonal.

To our knowledge, SeqWalk is the first orthogonal sequence design algorithm that is provably guaranteed to yield maximally sized SSM satisfying sequence libraries. SSM-satisfying sequence libraries designed by the partitioning of a Hamiltonian path (such as in SeqWalk) are maximally sized. This can be trivially proven by contradiction, by noting that every possible k-mer in the sequence space appears in the library. If there existed a larger library of SSM-satisfying sequences, it would use a larger number of k-mers, and thus would repeat k-mers, and not satisfy SSM. A more formal statement and proof can be found in Supplementary note 6. Building on fundamental results about de Bruijn graphs (11), we can obtain a closed form expression for the number of sequences in SeqWalk libraries under different design parameters. For alphabet size *m*, sequence length *L*, and SSM constraint *k*, the number of possible orthogonal sequences *N* is the number of nodes in the k-mer graph divided by the number of nodes required to represent a sequence of length *L*. More precisely,

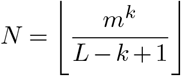

By solving the inverse problem given a desired library size of *N_d_*, sequence length *L*, and alphabet size *m*, we can choose the smallest *k* such that *N* ≥ *N_d_*. Designing a library using the resulting *k* value yields a maximally orthogonal library with the desired number of sequences.

Additionally, we can estimate the size of SeqWalk libraries after downstream filtering. For example, we can place lower bounds on the number of sequences present after a filtering for a specific sequence pattern of length *p* < *k*. The number of k-mers containing a specific pattern of length *p* is

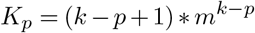

where *m* is the size of the alphabet. Since no k-mer appears in more than one sequence in the library, we must remove at most *K_p_* sequences from our library to remove all sequences containing a pattern of length *p*. As such, the size of the filtered library, *N_p_*, is

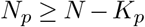

Such lower bounds are simple to determine for practically relevant pattern constraints, such as the prevention of homopolymeric regions (Fig S6). Additionally, we derive a lower bound on the size of SeqWalk libraries upon filtering for orthogonality with reverse complements (Supplemenary note 4). For the case of 3-letter libraries with odd *k*, we show that the size of a SeqWalk library that satisfies orthogonality with reverse complements, *N_rc_*, can be bounded by

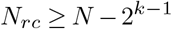

The size of SeqWalk libraries under GC content constraints is not as easily determined analytically. However, empirical results show that SeqWalk libraries have consistent distributions of GC content, resembling the binomial distribution expected of uniformly random sequences (Supplementary note 2). As such, these distributions can be used to estimate the size of SeqWalk libraries under GC content constraints.

### Implementation as a software tool

We have implemented the SeqWalk algorithm and additional filtering tools in a `pip` distributed Python package (`seqwalk`, source code available at github.com/storyetfall/seqwalk, documented at `seqwalk.readthedocs.io`). Additionally, we have developed an interactive, code-free, web-based SeqWalk interface in a publicly accessible Google Colaboratory notebook (link on `seqwalk.readthedocs.io`). We envision the use of `seqwalk` as a part of a sequence design pipeline, with downstream filtering (experimental validation, genomic homology filtering, etc.) as necessary for specific application contexts. Due to the simplicity of the underlying algorithms, we expect that others can implement our design method in other languages and development environments, and modify it as necessary.

## Discussion

We have presented SeqWalk, a method for efficiently producing maximally-sized orthogonal sequence libraries which are amenable to theoretical analyses. SeqWalk enables the design of orthogonal sequence libraries of unprecented size, with theoretical guarantees on maximality.

While SeqWalk is applicable to any orthogonal sequence design problem, the use of the SSM heuristic makes it more naturally applicable to certain kinds of design problems. In particular, SeqWalk is well suited for problems where nuanced biophysical properties (i.e. exact Δ*G*, strand displacement kinetics) need not be tightly controlled. We expect that SeqWalk will be valuable for the rapidly growing class of high-throughput biological methods that use synthetic DNA sequences as barcodes for different biomolecular features (i.e. samples, cells, protein targets, plasmids, etc.). Using SeqWalk to maximize the size of orthogonal sequence libraries can, in principle, increase the number of features that can be barcoded in such methods.

Additionally, the theoretical guarantees of SeqWalk libraries can be used to guide design choices in experimental method development. Using the results derived in this paper, one can understand the tradeoffs between design parameters and orthogonal sequence library size.

In the early 1990s, graph representations of biological sequences revolutionized the field of genomics, dramatically improving the quality and efficiency of de novo genome assembly (15). Since then, graph representations of sequences have become widespread as descriptive tools in bioinformatics, used to reconstruct naturally occurring biological sequences. In modern molecular biology and bioengineering, where the design of synthetic biological systems is fundamentally intertwined with the characterization of natural biological systems, there is growing interest in sequence representations amenable to design tasks (16, 17). However, outside of highly specialized applications (18, 19), graph representations of sequences are far less commonly used in design contexts. With SeqWalk, we demonstrate that graph-based sequence representations enable massive efficiency improvements in orthogonal sequence design. We are optimistic that graph representations of sequence space can similarly enable efficient solutions to other biological sequence design problems.

## Supporting information

Supplemental Notebooks 1-12

## ACKNOWLEDGEMENTS

We thank Jocelyn Kishi, Tatiana Brailovskaya, and Erik Winfree for thoughtful discussions. We thank Xiaokang Lun and Ninning Liu for feedback on the manuscript. We thank the Jupyter Project for maintaining open-source computational tools.

## Supplementary Note 1: Defining crosstalk

The words “orthogonality” and “crosstalk” are used frequently in molecular bioengineering, without very precise definitions. Here, we will try to be more precise about what we mean.

We consider two sequences (A, B) to have crosstalk if they can stably hybridize with each other’s reverse complements. In other words, if a complex between A and B*, or A* and B is likely to form, we consider A and B to have crosstalk.

If we think of A and B as probes, with A* and B* being their respective targets, we consider crosstalk to be the binding of a probe to an incorrect target. We do not by default consider binding between A and B to be crosstalk.

For many, but not all, applications, this is a sufficient characterization of crosstalk. In the case of multiplexed imaging, only a single probe (referred to as imager in the multiplexed imaging literature) is present in a sample at a given time (1). As such, we need not consider binding between probes. Analogously, in DNA similarity search, a single “query” probe is used to bind “target” strands, so binding between probe strands is not necessary.

In some applications, a stronger definition of crosstalk, including binding between probe strands, is necessary. For example, in single stranded tile self assembly, a pool of single strands, including “probes” and “targets” will be in well mixed solution. As such, binding between all strands must be considered crosstalk.

We consider this to be orthogonality including reverse complements, where A and B have crosstalk if any pair of A, A*, B, B* have significant binding (other than the desired A with A*, and B with B*). Sequence design under this stronger orthogonality constraint is discussed in the supplementary note on orthogonality with reverse complements, and the main text section on sequence design under additional constraints.

## Supplementary Note 2: GC content distributions in SeqWalk libraries

While we do not have a rigorous proof of this, we find empirically that the distributions of GC content in SeqWalk libraries is similar to that of totally random sequences. We expect, in a 4 letter alphabet, to have GC content that is binomially distributed with p=0.5 and n=L (where L is the sequence length). In a 3 letter alphabet (ACT or AGT) we expect to have GC content binomially distributed with 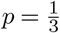 and *n* = *L.*

The variance of a binomial is *σ*^2^ (*n,p*) = *p* * (1 – *p*) * *n*, with *p* ∈ [0,1]. Since *p* * (1 – *p*) is globally maximized for *p* = 0.5, the variance of GC content is highest for the case of a 4 letter alphabet. This is in line with empirical results, shown below and in Supplemental File W.

**Fig S4.**
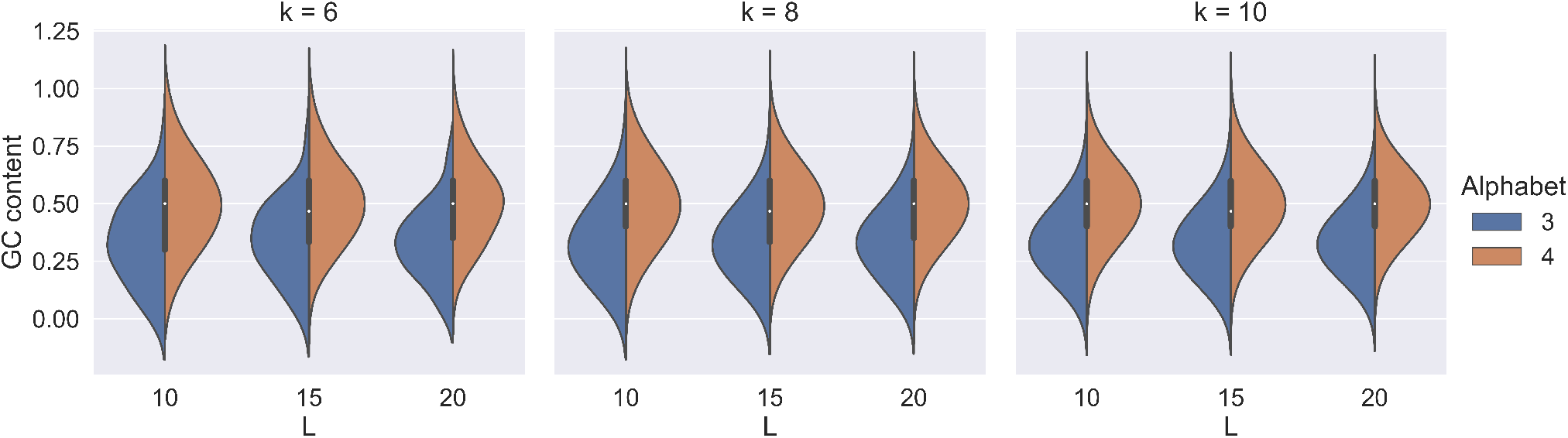
GC content distributions in various SeqWalk libraries. The 3 letter libraries are designed using {A, C, T}, while the 4 letter libraries use {A, C, T, G}. The code used to generate libraries can be found in Supplemental File X, the libraries can be found in Supplemental Files Y-Z, and the code used to make this plot can be found in Supplemental File W.

Since the GC content of 3 letter alphabet libraries is lower, a tighter window of GC content constraints can be used to obtain the same number of sequences (assuming that the extreme GC content sequences are those to be filtered out).

For filtering libraries for GC content, a naive algorithm, such as the one below, is efficient (constant time and memory in the size of the library).

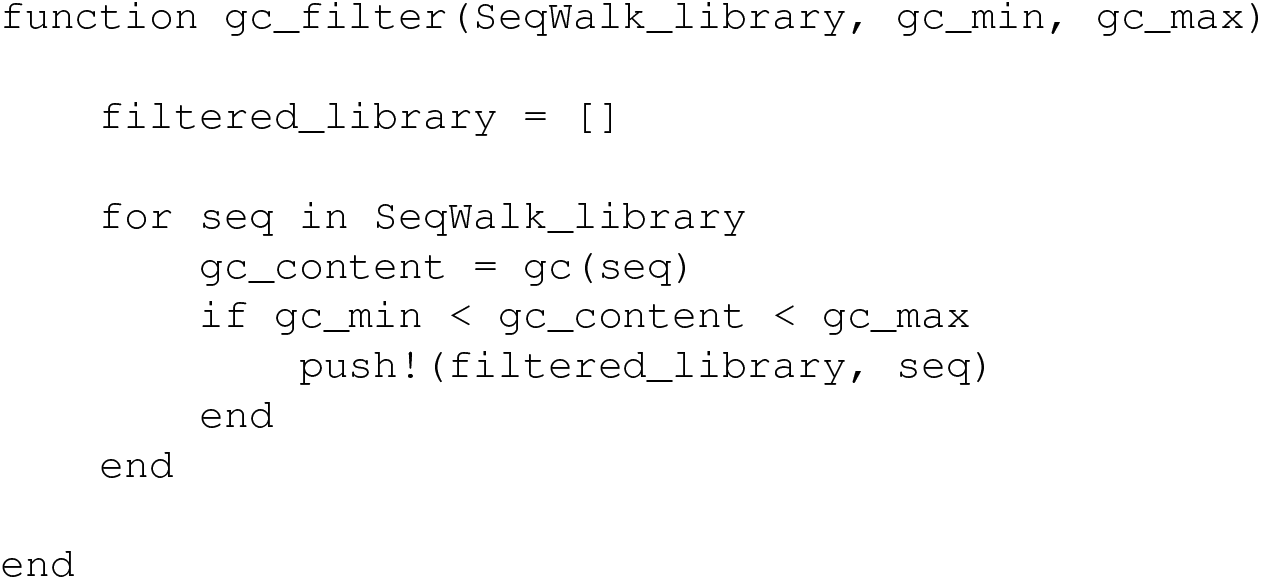

## Supplementary Note 3: Secondary structure in SeqWalk libraries

Empirically, we find that secondary structure is very uncommon in SeqWalk libraries constructed with {A, C, T} libraries. We use percentage of paired bases in the MFE structure as a measure of secondary structure prevalence in a sequence. We are aware that this is not an ideal measure based on (20), but we use it as it is simple to compute and relatively agnostic to experimental context.

**Fig S5.**
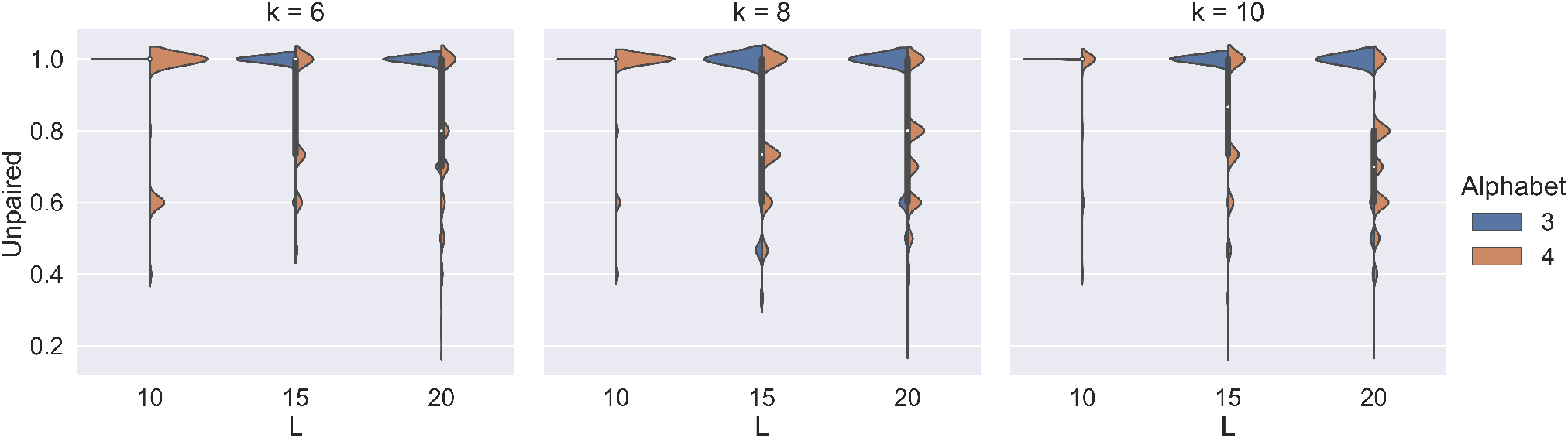
Distribution of fraction of unpaired bases in MFE structures in various SeqWalk libraries. An “Unpaired” value of 1 indicates no bound bases in the MFE structure of a sequence. The 3 letter libraries are designed using {A, C, T}, while the 4 letter libraries use {A, C, T, G}. The code used to generate libraries can be found in Supplemental File X, the libraries can be found in Supplemental Files Y-Z, and the code used to make this plot can be found in Supplemental File W.

To filter sequences for secondary structure, we can again use a naive algorithm, such as the one below, which is constant time and memory in the size of the library.

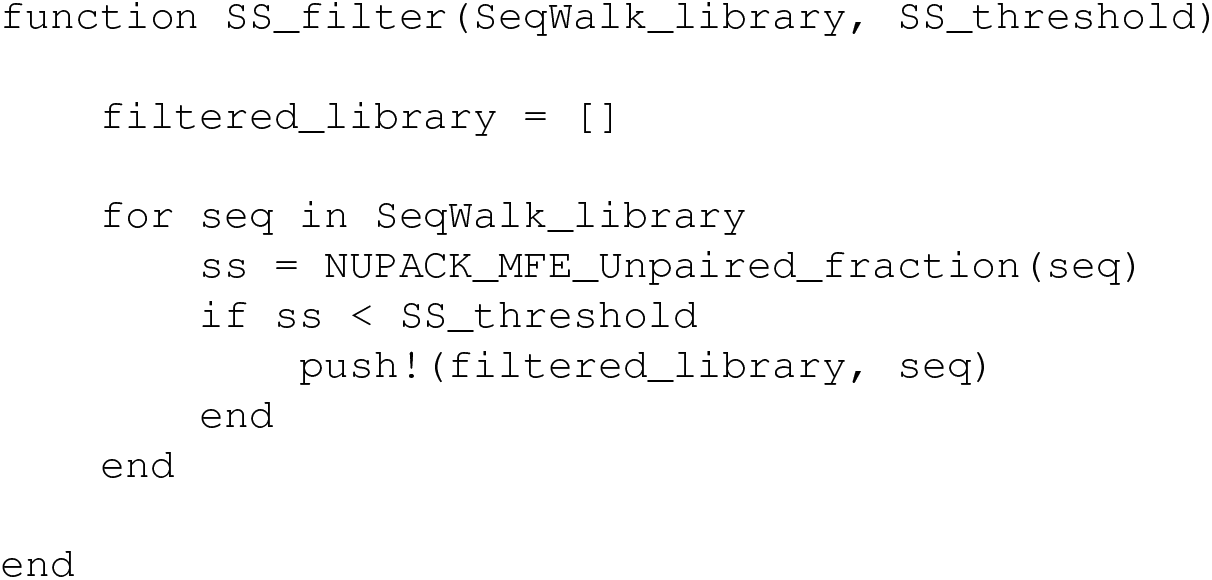

## Supplementary Note 4: Orthogonality with Reverse Complements

For the case of 3-letter alphabets and odd SSM *k* values, we present an efficient algorithm for selecting maximally sized orthogonal libraries which prevent crosstalk with reverse complements of sequences in the library. Without loss of generality, let’s consider the case of sequences constructed with an A, C, T library.

We seek to have a library which contains no repeated k-mers, and no k-mers whose reverse complement also appears in the library. K-mers containing “C” cannot have their reverse complements also appear in the library, since the library will not contain “G”. So, we only need to consider k-mers composed entirely of “A” and “T”.

In order to use k-mers whose reverse complement will not appear in the library, we seek partition all “AT” k-mers into two sets, such that the reverse complement of each sequence in a set appears in the other set.

In other words, if we consider a “Reverse Complement” graph, in which each node is a k-mer, and there is an edge between k-mers which are reverse complementary, we would like find a balanced bipartitioning of the graph.

Upon this partitioning, we can remove all sequences containing k-mers from one partition. Thus, the reverse complements of any k-mers that appear in the library will not be present.

For odd k, we can find such a partitioning by noting that the middle base in the k-mer will be different in its reverse complement. For example in a 5mer, the third base will never be the same as the third base of its reverse complement. As such, we can find a bipartition of the Reverse Complement graph by dividing the nodes into two sets, where all nodes in one set have “A” as the middle base, and all nodes in the other set have “T” as the middle base.

Below is the pseudocode for efficiently filtering sequences in a odd K, 3 letter alphabet library, such that orthogonality with reverse complements is respected.

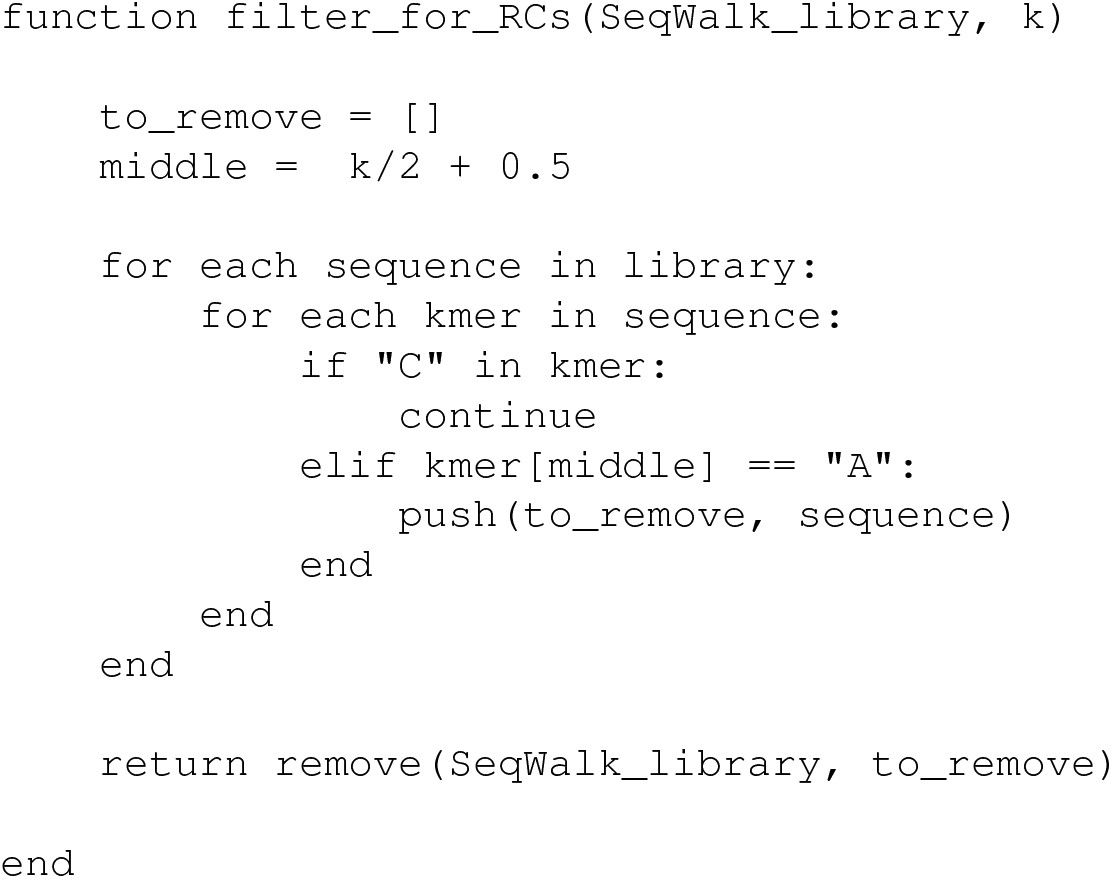

Using this algorithm, we can easily lower bound the size of a resulting library will be upon filtering. We know that there are 2^*k*^ kmers consisting entirely of A and T. Half of these k-mers will have “A” as the middle base. At most, we will remove one sequence from the libray for each kmer. As such, we can lowerbound the number of sequences upon reverse complementarity filtering, *Nrc* using

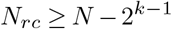

This theoretical result indicates that SeqWalk still produces relatively large sequence libraries upon such filtering. For example, for the case of 25nt barcodes with 3 letter code, SSM *k* = 13, and removal of reverse complements, we will have a sequence library with at least 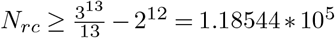 sequences.

In the case of a four-letter alphabet, filtering is an untenable solution because we cannot constrain reverse complementary k-mers to AT sequences. Instead, we use a modification of the Hierholzer algorithm, in which we mark both the visited k-mer and its reverse-complement “visited” during traversal. This method requires keeping track of visited nodes, and as such is less time/memory efficient than the shift rule traversal. Our implementation can be found in the `adapted_hierholzer` function in the `generation` module of the `seqwalk` source code (github.com/storyetfall/seqwalk)

## Supplementary Note 5: Preventing specific sequence patterns

We can place lower bounds on the number of sequences present after a filtering for a specific sequence pattern of length *p* < *k*. The number of k-mers containing a specific pattern of length *p* is

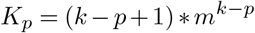

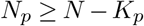

Such lower bounds are simple to determine for practically relevant pattern constraints, such as the prevention of homopolymeric regions.

For example, we can consider the case of preventing 4G regions, as well as 4N (any of 4A, 4T, 4C, 4G) regions. To lower bound the number of sequences after removing all 4N regions, we can use

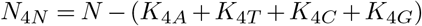

Below, we see plots of these bounds for various *k* and *L*.

**Fig S6.**
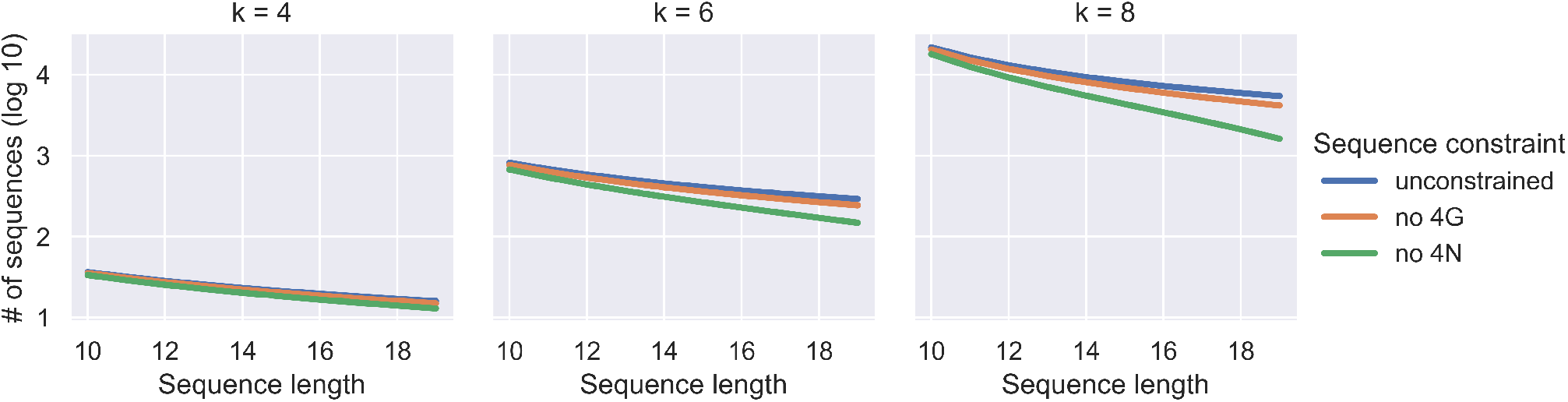
Lower bounds on library size for various design problems under different sequence pattern prevention constraints. In particular, we plot lower bounds for 4 letter SeqWalk libraries preventing 4G and preventing all 4N, in comparison to libraries with no pattern prevention constraints.

## Supplementary Note 6: Proof of maximality

### A. Definitions

- Sequence library: set of sequences of length *N* over alphabet of size *m*
- k-mer: subsequence of length *k*
- SSM satisfied for length *k*: no subsequence of length *k* appears more than once, for *k* < *N*
- Maximally sized SSM sequence library: A sequence library satisfying SSM for length *k* with size such that no larger sequence library satisfying SSM for length *k* exists.

### B. Lemma 1.

A maximally sized sequence library that satisfies SSM for length k contains at most *m^k^* distinct k-mers.

### C. Proof of Lemma 1.

Assume for the sake of contradiction that there exists an SSM satisfying library for length *k*, which has *K* > *m^k^* k-mers. Since there are only *m^k^* possible k-mers, by the pigeonhole principle, at least one k-mer must appear > 1 times in the library. Since a k-mer appears more than once in the library, it does not satisfy SSM. We have arrived a contradiction.

### D. Theorem 1.

A sequence library generated by the partitioning of a Hamiltonian path in a *k* de Bruijn graph is a maximally sized SSM sequence library for length *k*.

### E. Proof of Theorem 1.

By definition, the number of k-mers in such a library is equal to the number of nodes in the corresponding de Bruijn graph. The number of nodes in the de Bruijn graph, by definition, is *m^k^*. By Lemma 1, a maximally sized sequence library that satisfies SSM for length contains at most *m^k^* k-mers. Thus, no larger SSM satisfying library exists.

## Bibliography

1. Sinem K Saka, Yu Wang, Jocelyn Y Kishi, Allen Zhu, Yitian Zeng, Wenxin Xie, Koray Kirli, Clarence Yapp, Marcelo Cicconet, Brian J Beliveau, Sylvain W Lapan, Siyuan Yin, Millicent Lin, Edward S Boyden, Pascal S Kaeser, German Pihan, George M Church, and Peng Yin. Immuno-SABER enables highly multiplexed and amplified protein imaging in tissues. Nat. Bìotechnol., 37(9):1080–1090, September 2019.

2. Allon M Klein, Linas Mazutis, Ilke Akartuna, Naren Tallapragada, Adrian Veres, Victor Li, Leonid Peshkin, David A Weitz, and Marc W Kirschner. Droplet barcoding for single-cell transcriptomics applied to embryonic stem cells. Cell, 161(5):1187–1201, May 2015.

3. Z J Gartner and D R Liu. The generality of DNA-templated synthesis as a basis for evolving non-natural small molecules. J. Am. Chem. Soc., 123(28):6961–6963, July 2001.

4. Arturo Casini, Georgia Christodoulou, Paul S Freemont, Geoff S Baldwin, Tom Ellis, and James T MacDonald. R2oDNA designer: computational design of biologically neutral synthetic DNA sequences. ACS Synth. Biol., 3(8):525–528, August 2014.

5. Timothy C Yu, Winnie L Liu, Marcia S Brinck, Jessica E Davis, Jeremy Shek, Grace Bower, Tal Einav, Kimberly D Insigne, Rob Phillips, Sriram Kosuri, and Guillaume Urtecho. Multiplexed characterization of rationally designed promoter architectures deconstructs combinatorial logic for IPTG-inducible systems. Nat. Commun., 12(1):325, January 2021.

6. Qikai Xu, Michael R Schlabach, Gregory J Hannon, and Stephen J Elledge. Design of 240,000 orthogonal 25mer DNA barcode probes. Proc. Natl. Acad. Sci. U. S. A., 106(7): 289–2294, February 2009.

7. A Marathe, A E Condon, and R M Corn. On combinatorial DNA word design. J. Comput. Biol.,8(3):201–219, 2001.

8. Constantine G Evans and Erik Winfree. DNA sticky end design and assignment for robust algorithmic self-assembly. In DNA Computing and Molecular Programming, pages 61–75. Springer International Publishing, 2013.

9. N C Seeman. De novo design of sequences for nucleic acid structural engineering. J. Biomol. Struct. Dyn., 8(3):573–581, December 1990.

10. D D Shoemaker, D A Lashkari, D Morris, M Mittmann, and R W Davis. Quantitative phenotypic analysis of yeast deletion mutants using a highly parallel molecular bar-coding strategy Nat. Genet., 14(4):450–456, December 1996.

11. T van Aardenne-Ehrenfest and Nicolaas Govert de Bruijn. Circuits and trees in oriented linear graphs. In Classic papers in combinatorics, pages 149–163. Springer, 2009.

12. Joe Sawada, Aaron Williams, and Dennis Wong. A surprisingly simple de bruijn sequence construction. Discrete Math., 339(1):127–131, January 2016.

13. Joe Sawada, Aaron Williams, and Dennis Wong. A simple shift rule for k-ary de bruijn sequences. Discrete Math., 340(3):524–531, March 2017.

14. Hierholzer and Wiener. Über die möglichkeit, einen linienzug ohne wiederholung und ohne unterbrechung zu umfahren. Math. Ann., 1873.

15. R M Idury and M S Waterman. A new algorithm for DNA sequence assembly. J. Comput. Biol., 2(2):291–306, 1995.

16. Johannes Linder and Georg Seelig. Fast differentiable DNA and protein sequence optimization for molecular design. May 2020.

17. Eli N Weinstein, Alan N Amin, Will Grathwohl, Daniel Kassler, Jean Disset, and Debora S Marks. Optimal design of stochastic DNA synthesis protocols based on generative sequence models. October 2021.

18. S Roweis, E Winfree, R Burgoyne, N V Chelyapov, M F Goodman, P W Rothemund, and L M Adleman. A sticker-based model for DNA computation. J. Comput. Biol., 5(4):615–629, 1998.

19. Yaron Orenstein and Bonnie Berger. Efficient design of compact unstructured RNA libraries covering all k-mers. J. Comput. Biol., 23(2):67–79, February 2016.

20. Robert M Dirks, Milo Lin, Erik Winfree, and Niles A Pierce. Paradigms for computational nucleic acid design. Nucleic Acids Res., 32(4):1392–1403, February 2004.

