## Supplemental Notebooks 1-12 for "Efficient algorithms for designing maximally sized orthogonal DNA sequence libraries": SF2_performance_and_constraints_plots.ipynb

```
# %% [code]
import pandas as pd
import seaborn as sns
import numpy as np
import matplotlib.pyplot as plt

# %% [code]
perf = pd.read_csv("perf.csv")

a_t = {"pairwise": "pairwise", "deBruijn" : "SeqWalk"}
perf["Algorithm"] = [a_t[s] for s in perf["algorithm"]]
perf["# of sequences"] = perf["sequences"]
perf["Time (s)"] = perf["time"] / 10**6

perf["time"] = perf["time"] / 10**6

# %% [code]
sns.set(font_scale=1.6)
g = sns.FacetGrid(perf, row="k", col="L", hue="Algorithm", sharex=False, sharey=False)
g.map_dataframe(sns.lineplot, x = "Time (s)", y= "# of sequences", lw=4)
g.add_legend()
g.figure.savefig("Performance.png", dpi=600)

# %% [code]
# MAX SSM LIBRARY SIZE PLOT

def f(L, k):
    return 4**k / (1+L-k)

def f_p4G(L, k):
    return f(L, k) - ((k-4+1) * 4**(k-4))
    
def f_p4N(L, k):
    return f(L, k) - 3*((k-4+1) * 4**(k-4))


ks = [4, 6, 8]
Ls = range(10, 20)

d = {
    
    "k": [],
    "Length": [],
    "# of sequences (log 10)": [],
    "Sequence constraint": []
}

for k in ks:
    for L in Ls:
        
        d["k"].append(k)
        d["Length"].append(L)
        d["# of sequences (log 10)"].append(np.log10(f(L, k)))
        d["Sequence constraint"].append("unconstrained")
        
        d["k"].append(k)
        d["Length"].append(L)
        d["# of sequences (log 10)"].append(np.log10(f_p4G(L, k)))
        d["Sequence constraint"].append("no 4G")
        
        d["k"].append(k)
        d["Length"].append(L)
        d["# of sequences (log 10)"].append(np.log10(f_p4N(L, k)))
        d["Sequence constraint"].append("no 4N")


d["Sequence length"] = d["Length"]
        
hm = pd.DataFrame(d)

# %% [code]
# def draw_heatmap(*args, **kwargs):
#     data = kwargs.pop('data')
#     d = data.pivot(index=args[1], columns=args[0], values=args[2])
#     sns.heatmap(d, vmin = 1, vmax=6, **kwargs)

hg = sns.FacetGrid(hm, col="k", hue="Sequence constraint")
hg.map_dataframe(sns.lineplot, "Sequence length", "# of sequences (log 10)", lw=3)
hg.add_legend()
hg.figure.savefig("Constraints.png", dpi=600)

# %% [code]


# %% [code]
```
