## Supplemental Notebooks 1-12 for "Efficient algorithms for designing maximally sized orthogonal DNA sequence libraries": SF3_library_generation.ipynb

```
# %% [code]
using Random
alph = ["A", "C", "T", "G"];

# %% [code]
"""
    isNecklace(seq)

## summary
computes if a sequence is a necklace, as defined by Wong 2017. 

## scaling
O(n)

## usage notes
works for strings, lists, numbers, bools
"""
function isNecklace(seq)
    
    p = 1
    
    for i in 2:length(seq)
        if seq[i-p] < seq[i]
            p = i
        elseif seq[i-p] > seq[i]
            return false
        end
    end
    
    if mod(length(seq), p) == 0
        return true
    else
        return false
    end
    
end


"""
    simpleShift(n, k)

## summary
executes simpleShift algorithm presented in Wong 2017.

## scaling
O(nk)
"""
function simpleShift(n, k)
    
    seq = rand(1:k, n)

    while true
        push!(seq, f(seq[end-n+1:end], k))
        if seq[end-n+1:end] == seq[1:n]
            break
        end
    end
    
    seq
end

"""
    f(seq, k)

## summary
helper function f presented in Wong 2017

## scaling
O(n)

## usage notes
requires list of ints
"""
function f(seq::Array, k)
    p = 1
    if seq[1] == k
        if sum(seq[2:end]) == length(seq) - 1
            return 1
        end
        for i in 0:k-2
            if !isNecklace([seq[2:end] ; k-i])
                return k-i
            end
        end
        return 1
    elseif isNecklace([seq[2:end] ; mod(seq[1], k) + 1])
        return mod(seq[1], k) + 1
    else
        return seq[1]
    end
end

"""
    partitionCycle(simpleShift(3, 3), 3, 4)

## summary
split cycle into sequences of length L

## scaling
O(n/(L-k))
"""
function partitionCycle(cycle, k, L)
    library = []
    for i in 1:L-k+1:length(cycle)-L
        seq = []
        for j in 0:L-1
            push!(seq, cycle[i+j])
        end
        push!(library, seq)
    end
    library
end

function intToDNA(seq)
    join([alph[i] for i in seq])
end

# %% [code]
alphabet_size = 4 # number of allowable bases

for SSM_k in [6, 8, 10]
    for seq_L in [10, 15, 25]

        @time library = partitionCycle(simpleShift(SSM_k, alphabet_size), SSM_k, seq_L)
        seqs = intToDNA.(library)


        filename = "SSM_$SSM_k-L_$(seq_L)-a_$(alphabet_size)" # file to write sequences to

        shuffle!(seqs)

        open(filename, "w+") do f
            for seq in seqs
                write(f, "$seq\n")
            end
        end
    end
end

# %% [code]
alphabet_size = 3 # number of allowable bases

for SSM_k in [6, 8, 10]
    for seq_L in [10, 15, 25]

        @time library = partitionCycle(simpleShift(SSM_k, alphabet_size), SSM_k, seq_L)
        seqs = intToDNA.(library)


        filename = "SSM_$SSM_k-L_$(seq_L)-a_$(alphabet_size)" # file to write sequences to

        shuffle!(seqs)

        open(filename, "w+") do f
            for seq in seqs
                write(f, "$seq\n")
            end
        end
    end
end

# %% [code]
```
