## Supplemental Notebooks 1-12 for "Efficient algorithms for designing maximally sized orthogonal DNA sequence libraries": SF6_supplement_gc.ipynb

````
# %% [markdown]
# Supplement: GC Content

While we do not have a rigorous proof of this, we find empirically that the distributions of GC content in SeqWalk libraries is similar to that of totally random sequences. We expect, in a 4 letter alphabet, to have GC content that is binomially distributed with p=0.5 and n=L (where L is the sequence length). In a 3 letter alphabet (ACT or AGT) we expect to have GC content binomially distributed with $p = \frac{1}{3}$ and $n=L$.

Since the variance of a binomial is $\sigma^2 (n, p) = p * (1 - p) * n$. Since $p*(1-p)$ is globally maximized for $p=0.5$, the variance of GC content is highest for the case of a 4 letter alphabet. This is in line with empirical results, shown below.

Since the GC content of 3 letter alphabet libraries is lower, a tighter window of GC content constraints can be used to obtain the same number of sequences (assuming that the extreme GC content sequences are those to be filtered out).

For filtering libraries for GC content, a naive algorithm is efficient.
```
function gc_filter(SeqWalk_library, gc_min, gc_max)
    filtered_library = []
    
    for seq in SeqWalk_library
        gc_content = gc(seq)
        if gc_min < gc_content < gc_max
            push!(filtered_library, seq)
        end
    end
end

```

# %% [code]
import nupack as nu
import numpy as np
import os
import matplotlib.pyplot as plt
import seaborn as sns
import pandas as pd

# %% [code]
def gc(seq):
    return sum( 1 for s in seq if s in ["G", "C"])/len(seq)

# %% [code]
d = {
    "Alphabet" : [],
    "k" : [],
    "L" : [],
    "GC content" : []
}

for SSM_k in [6, 8, 10]:
    for L in [10, 15, 20]:
        for a in [3, 4]:
            filename = "SSM_%d-L_%d-a_%d" % (SSM_k, L, a)
            f = open(filename, "r")
            seqs = [s.strip() for s in f.readlines()]
            f.close
            for seq in seqs:
                d["Alphabet"].append(a)
                d["k"].append(SSM_k)
                d["L"].append(L)
                d["GC content"].append(gc(seq))

data = pd.DataFrame(d)

# %% [code]
sns.set(font_scale=1.6)
# g = sns.FacetGrid(data, row="k", col="L", hue="Alphabet")
g = sns.catplot(data = data, y = "GC content", x="L", col="k", hue="Alphabet", 
                split=True, kind="violin", bw=0.5)
g.figure.savefig("GC.png", dpi=600)

# %% [code]
var = data.groupby("Alphabet").var(ddof=1)["GC content"]

print("Variance of 3 letter: %f" % var[3])
print("Variance of 4 letter: %f" % var[4])

# %% [code]
g.figure.savefig("GC.png", dpi=600)

# %% [code]
````
