## Supplemental Notebooks 1-12 for "Efficient algorithms for designing maximally sized orthogonal DNA sequence libraries": SF9_supplement_performance_comparison.ipynb

```
# %% [code]
using Random
using Dates
using CSV
using DataFrames

# %% [code]
function pairwise_comparison(L, k)
    lib = []
    kmers = []
    times = [time_ns()*1e-3]
    iter = 0
    while true
        cand_seq = randstring("ACTG", L)
        ssm = true
        iter += 1
        if iter > 10
#             println("iter > 1")
#             println(Dates.value(Dates.now())- times[1])
            if time_ns()*1e-3 - times[1] > 1 * 10^6
                return times
            end
        end
        for i in 1:L-k+1
            if cand_seq[i:i+k-1] in kmers
                ssm = false
                break
            end
        end
        if ssm
            iter = 0
            push!(lib, cand_seq)
#             println(times)
            push!(times, time_ns()*1e-3) # microseconds
            for i in 1:L-k+1
                push!(kmers, cand_seq[i:i+k-1])
            end
        end
    end
    
end

# %% [code]
alph = ["A", "C", "G", "T"]
"""
    isNecklace(seq)

## summary
computes if a sequence is a necklace, as defined by Wong 2017. 

## scaling
O(n)

## usage notes
works for strings, lists, numbers, bools
"""
function isNecklace(seq)
    
    p = 1
    
    for i in 2:length(seq)
        if seq[i-p] < seq[i]
            p = i
        elseif seq[i-p] > seq[i]
            return false
        end
    end
    
    if mod(length(seq), p) == 0
        return true
    else
        return false
    end
    
end


"""
    simpleShift(n, k)

## summary
executes simpleShift algorithm presented in Wong 2017.

## scaling
O(nk)
"""
function simpleShift(n, k)
    
    seq = rand(1:k, n)

    while true
        push!(seq, f(seq[end-n+1:end], k))
        if seq[end-n+1:end] == seq[1:n]
            break
        end
    end
    
    seq
end

"""
    f(seq, k)

## summary
helper function f presented in Wong 2017

## scaling
O(n)

## usage notes
requires list of ints
"""
function f(seq::Array, k)
    p = 1
    if seq[1] == k
        if sum(seq[2:end]) == length(seq) - 1
            return 1
        end
        for i in 0:k-2
            if !isNecklace([seq[2:end] ; k-i])
                return k-i
            end
        end
        return 1
    elseif isNecklace([seq[2:end] ; mod(seq[1], k) + 1])
        return mod(seq[1], k) + 1
    else
        return seq[1]
    end
end

"""
    partitionCycle(simpleShift(3, 3), 3, 4)

## summary
split cycle into sequences of length L

## scaling
O(n/(L-k))
"""
function partitionCycle(cycle, k, L)
    library = []
    for i in 1:L-k+1:length(cycle)-L
        seq = []
        for j in 0:L-1
            push!(seq, cycle[i+j])
        end
        push!(library, seq)
    end
    library
end

function intToDNA(seq)
    join([alph[i] for i in seq])
end

# %% [code]
n = 10
ks = []
Ls = []
algs = []
ts = []
seqs = []

for k in [4, 6, 8]
    for L in [10, 15, 20]        
        
        trials = []
        for i in 1:n
            push!(trials, pairwise_comparison(L, k))
        end
        
        print("pairwise done")
        
        m = minimum(length.(trials))

        t = [sum([trials[j][i] - trials[j][1] for j in 1:n]) for i in 1:m]
        
        
        ts = vcat(ts, t)
        Ls = vcat(Ls, [L for i in 1:length(t)])
        ks = vcat(ks, [k for i in 1:length(t)])
        algs = vcat(algs, ["pairwise" for i in 1:length(t)])
        seqs = vcat(seqs, collect(0:length(t)-1))
        
        start = time_ns()
        res = nothing
        for i in 1:n
            library = partitionCycle(simpleShift(k, 4), k, L)
            res = intToDNA.(library)
        end
        elapsed = ((time_ns() - start)*1e-3)/n
        
        ts = vcat(ts, [0, elapsed])
        ks = vcat(ks, [k, k])
        Ls = vcat(Ls, [L, L])
        algs = vcat(algs, ["deBruijn", "deBruijn"])
        seqs = vcat(seqs, [0, length(res)])
    end
end

out = DataFrame(k = ks, L = Ls, algorithm = algs, time = ts, sequences = seqs)

# %% [code]
CSV.write("perf.csv", out)

# %% [code]
```
