## Supplemental Notebooks 1-12 for "Efficient algorithms for designing maximally sized orthogonal DNA sequence libraries": SF10_supplement_secondary_struct.ipynb

```
# %% [code]
import nupack as nu
import numpy as np
import os
import matplotlib.pyplot as plt
import seaborn as sns
import pandas as pd

c = nu.Model(material="dna", celsius=22, sodium=0.05, magnesium=0.002)

# %% [code]
def mfe_unpaired(seq, model):
    A = nu.Strand(seq, name="A")
    return sum(1 for s in str(nu.mfe([A], model=model)[0].structure) if s == ".")/len(seq)

def gc(seq):
    return sum( 1 for s in seq if s in ["G", "C"])/len(seq)

# %% [code]
d = {
    "Alphabet" : [],
    "k" : [],
    "L" : [],
    "Unpaired" : [],
    "GC content" : []
}

for SSM_k in [6, 8, 10]:
    for L in [10, 15, 20]:
        for a in [3, 4]:
            filename = "SSM_%d-L_%d-a_%d" % (SSM_k, L, a)
            f = open(filename, "r")
            seqs = [s.strip() for s in f.readlines()]
            f.close
            for seq in seqs:
                d["Alphabet"].append(a)
                d["k"].append(SSM_k)
                d["L"].append(L)
                d["Unpaired"].append(mfe_unpaired(seq, c))
                d["GC content"].append(gc(seq))

data = pd.DataFrame(d)
data.to_csv("secondary_structs.csv")

# %% [code]
sns.set(font_scale=1.6)
# g = sns.FacetGrid(data, row="k", col="L", hue="Alphabet")
g = sns.catplot(data = data, y = "Unpaired", x="L", col="k", hue="Alphabet", 
                split=True, kind="violin", bw=0.1)
g.figure.savefig("Unpaired.png", dpi=600)

# %% [code]


# %% [code]


# %% [code]
```
