## Supplemental Notebooks 1-12 for "Efficient algorithms for designing maximally sized orthogonal DNA sequence libraries": SF12_software_demo.ipynb

```
# %% [code]
from seqwalk import design
from seqwalk import analysis
import seaborn as sns
import matplotlib.pyplot as plt
import random

# %% [code]
design.max_orthogonality(200, # number of sequences
                         25,  # sequence length
                         # ... other design constraints
                        )

# %% [code]
small_25mer_library = design.max_orthogonality(200, 25, 
                                               alphabet="ACGT", prevented_patterns=[])

# %% [code]
h_crosstalk = analysis.hamming_matrix(small_25mer_library)

p = sns.heatmap(h_crosstalk, xticklabels=False, yticklabels=False)
cbar = p.collections[0].colorbar
cbar.ax.tick_params(labelsize=15)
#plt.title("Hamming distance matrix")
plt.show()

# %% [code]
[random.choice(small_25mer_library) for i in range(10)]

# %% [code]
```
