## Supplemental Notebooks 1-12 for "Efficient algorithms for designing maximally sized orthogonal DNA sequence libraries": SF_1_SeqWalk.ipynb

```
# %% [markdown]
# SeqWalk orthogonal sequence library generation

Using this notebook, you can generate orthogonal sequences with the computational method described in \cite{preprint}. Simply input design parameters in text boxes -- no code necessary!

*Built from [Julia Colab Notebook Template](https://colab.research.google.com/github/ageron/julia_notebooks/blob/master/Julia_Colab_Notebook_Template.ipynb)*

## Instructions

1. Copy this notebook: _File_ > _Save a copy in Drive_ (you will need a Google account). 
2. Execute the following "setup" cell (click the play button). This will take a couple of minutes.
3. Reload this page (press Ctrl+R or equivalent).
4. Execute the next "load" cell. This will take a few seconds.
5. Fill out the form in the last cell with desired design parameters, and run! 
6. Navigate to the Files tab (left panel) and download the output text file with sequences.

(Note: Steps 1-4 need to be done only once per instance. Steps 5 and 6 can be repeated with different parameters, without repeating 1-4.)

# %% [code]
#@title Set up the notebook
%%shell
set -e

#---------------------------------------------------#
JULIA_VERSION="1.7.1" # any version ≥ 0.7.0
JULIA_PACKAGES="IJulia Random"
JULIA_PACKAGES_IF_GPU="CUDA" # or CuArrays for older Julia versions
JULIA_NUM_THREADS=2
#---------------------------------------------------#

if [ -n "$COLAB_GPU" ] && [ -z `which julia` ]; then
  # Install Julia
  JULIA_VER=`cut -d '.' -f -2 <<< "$JULIA_VERSION"`
  echo "Installing Julia $JULIA_VERSION on the current Colab Runtime..."
  BASE_URL="https://julialang-s3.julialang.org/bin/linux/x64"
  URL="$BASE_URL/$JULIA_VER/julia-$JULIA_VERSION-linux-x86_64.tar.gz"
  wget -nv $URL -O /tmp/julia.tar.gz # -nv means "not verbose"
  tar -x -f /tmp/julia.tar.gz -C /usr/local --strip-components 1
  rm /tmp/julia.tar.gz

  # Install Packages
  for PKG in `echo $JULIA_PACKAGES`; do
    echo "Installing Julia package $PKG..."
    julia -e 'using Pkg; pkg"add '$PKG'; precompile;"' &> /dev/null
  done

  # Install kernel and rename it to "julia"
  echo "Installing IJulia kernel..."
  julia -e 'using IJulia; IJulia.installkernel("julia", env=Dict(
      "JULIA_NUM_THREADS"=>"'"$JULIA_NUM_THREADS"'"))'
  KERNEL_DIR=`julia -e "using IJulia; print(IJulia.kerneldir())"`
  KERNEL_NAME=`ls -d "$KERNEL_DIR"/julia*`
  mv -f $KERNEL_NAME "$KERNEL_DIR"/julia  

  echo ''
  echo "Successfully installed `julia -v`!"
  echo "Please reload this page (press Ctrl+R, ⌘+R, or the F5 key)"
fi

# %% [code]
#@title Load design code

using Random
using DelimitedFiles

"""
    isNecklace(seq)

## summary
computes if a sequence is a necklace, as defined by Wong 2017. 

## scaling
O(n)

## usage notes
works for strings, lists, numbers, bools
"""
function isNecklace(seq)
    
    p = 1
    
    for i in 2:length(seq)
        if seq[i-p] < seq[i]
            p = i
        elseif seq[i-p] > seq[i]
            return false
        end
    end
    
    if mod(length(seq), p) == 0
        return true
    else
        return false
    end
    
end


"""
    simpleShift(n, k)

## summary
executes simpleShift algorithm presented in Wong 2017.

## scaling
O(nk)
"""
function simpleShift(n, k)
    
    seq = rand(1:k, n)

    while true
        push!(seq, f(seq[end-n+1:end], k))
        if seq[end-n+1:end] == seq[1:n]
            break
        end
    end
    
    seq
end

"""
    f(seq, k)

## summary
helper function f presented in Wong 2017

## scaling
O(n)

## usage notes
requires list of ints
"""
function f(seq::Array, k)
    p = 1
    if seq[1] == k
        if sum(seq[2:end]) == length(seq) - 1
            return 1
        end
        for i in 0:k-2
            if !isNecklace([seq[2:end] ; k-i])
                return k-i
            end
        end
        return 1
    elseif isNecklace([seq[2:end] ; mod(seq[1], k) + 1])
        return mod(seq[1], k) + 1
    else
        return seq[1]
    end
end

"""
    partitionCycle(simpleShift(3, 3), 3, 4)

## summary
split cycle into sequences of length L

## scaling
O(n/(L-k))
"""
function partitionCycle(cycle, k, L)
    library = []
    for i in 1:L-k+1:length(cycle)-L
        seq = []
        for j in 0:L-1
            push!(seq, cycle[i+j])
        end
        push!(library, seq)
    end
    library
end

function intToDNA(seq, alphabet)
    join([alphabet[i] for i in seq])
end;

println("Successfully loaded design code :)")


# %% [code]
#@title # Design sequence library { vertical-output: true, display-mode: "form" }

#@markdown ## Input parameters
#@markdown ### 1. Alphabet symbols (separated by commas) 
alphabet = "A,C,G,T" #@param {type:"string"}
#@markdown ### 2. SSM k value (length of unrepeated substrings)  
SSM_k =  12#@param {type:"integer"}
#@markdown ### 3. Length of sequences  
seq_L =  25#@param {type:"integer"}
#@markdown ### 4. Filename to save sequences
filename = "sequences.txt" #@param {type:"string"}
#@markdown ## Now run the cell to design sequences!

@assert !(seq_L < SSM_k)

alphabet = split(alphabet, ",") # allowable letters in sequence

@time library = partitionCycle(simpleShift(SSM_k, length(alphabet)), 
                                            SSM_k, seq_L)
seqs = [intToDNA(seq, alphabet) for seq in library]

writedlm(filename, seqs, '\n')

# %% [code]
```
